# Characterization and description of *Faecalibacterium butyricigenerans* sp. nov. and *F. longum* sp. nov., isolated from human faeces

**DOI:** 10.1101/2020.12.09.414284

**Authors:** Yuanqiang Zou, Xiaoqian Lin, Wenbin Xue, Ying Dai, Karsten Kristiansen, Liang Xiao

## Abstract

Exploiting a pure culture strategy to investigate the composition of human gut microbiota, two novel anaerobes, designated strains AF52-21^T^ and CM04-06^T^, were isolated from faeces of two healthy Chinese donors and characterized using a polyphasic approach. The two strains were Gram-stain-negative, non-motile, and rod-shaped. Both strains grew optimally at 37°C and pH 7.0. Phylogenetic analysis based on 16S rRNA gene sequences revealed that the two strain clustered with species of the genus *Faecalibacterium* and were most closely related to *Faecalibacterium prausnitzii* ATCC 27768^T^ with sequence similarity of 97.18% and 96.87%, respectively. The two isolates shared a 16S rRNA gene sequence identity of 98.69%. Draft genome sequencing was performed for strains AF52-21^T^ and CM04-06^T^, generating genome sizes of 2.85 Mbp and 3.01 Mbp. The calculated average nucleotide identity values between the genomes of the strains AF52-21^T^ and CM04-06^T^ compared to *Faecalibacterium prausnitzii* ATCC 27768^T^ were 83.20% and 82.54%, respectively, and 90.09% when comparing AF52-21^T^ and CM04-06^T^. Both values were below the previously proposed species threshold (95%), supporting their recognition as novel species in the genus *Faecalibacterium*. The genomic DNA G+C contents of strain AF52-21^T^ and CM04-06^T^ calculated from genome sequences were 57.77 mol% and 57.51 mol%, respectively. Based on the phenotypic, chemotaxonomic and phylogenetic characteristics, we conclude that both strains represent two new *Faecalibacterium* species, for which the names *Faecalibacterium butyricigenerans* sp. nov. (type strain AF52-21^T^ = CGMCC 1.5206^T^ = DSM 103434^T^) and *Faecalibacterium longum* sp. nov. (type strain CM04-06^T^ = CGMCC 1.5208^T^ = DSM 103432^T^) are proposed.

## Introduction

The human gastrointestinal^1^ tract harbours complex microbial communities^2^, dominated by bacteria from the phyla *Bacteroidetes* and *Firmicutes*^3-6^. The composition and diversity of the gut microbiota are affected by numerous factors, including host genetics^7^, long-term diet^8,9^, drugs^1,10,11^ and several other environmental factors^12^. Evidence suggests that the composition of the microbiota is associated with the development of obesity^4,13-15^, diabetes^16,17^, inflammatory bowel disease^18,19^, colorectal cancer^20,21^, and non-alcoholic fatty liver disease^22,23^. Therefore, the composition and function of the microbial species living in our gut are crucial importance for maintenance of health. Short-chain fatty acids (SCFAs), produced by fermentation of dietary fibre by several abundant genera of the intestinal microbiota, including *Roseburia, Eubacterium* and *Faecalibacterium*^24^, have been reported to elicit beneficial effects on energy metabolism and for prevention of colonization of pathogens^25^. The genus *Faecalibacterium* as an abundant butyric acid-producing bacterium colonizing the human gut displays anti-inflammatory effects and may be used as a potential probiotics for treatment of gut inflammation^26,27^.

The genus *Faecalibacterium*, belonging to the family *Ruminococcaceae* within the order *Clostridiales*, comprises only one validated species, *Faecalibacterium prausnitzii*^28^, and two non-validly published species, *Faecalibacterium moorei*^29^ and *Faecalibacterium hominis*^30^, all originally isolated from human faeces. *F. prausnitzii* is a gram-negative non-spore-forming and strictly anaerobic rod-shaped bacterium. The genomic G+C content of genus *Faecalibacterium* ranges from 47% to 57%^31^. The fermentation products from glucose are butyrate, D-lactate and formate. In the present study, we describe two novel species of the genus *Faecalibacterium* by using polyphasic taxonomy along with whole genome sequence analysis.

## Results and discussion

### Phenotypic and Chemotaxonomic Characterization

Both strains (AF52-21^T^ and CM04-06^T^) were obligate anaerobic, Gram-stain-negative, non-spore-forming, non-motile and rod-shaped bacteria **(Fig. 1)**. After incubation on MPYG agar at 37°C for 2 days, the colonies appeared 1.0-2.0 mm in diameter, round, creamy white to yellowish, convex and opaque with entire margins for AF52-21^T^ and 2.0 mm in diameter, round, yellowish, slightly convex and opaque with entire margins for CM04-06^T^. The growth temperature was 20-42°C (optimum 37°C) for AF52-21^T^ and 30-45°C (optimum 37°C) for CM04-06^T^. Growth was observed at pH 6.0-7.5 (optimum 7.0-7.5) for AF52-21^T^ and pH 5.0-8.0 (optimum 7.0-7.5) for CM04-06^T^. Strains AF52-21^T^ and CM04-06^T^ grew with 0-1% and 0-3% NaCl, respectively. Both strains were catalase-negative. The major metabolic end products for strains AF52-21^T^ and CM04-06^T^ were acetic acid, formic acid, butyric acid and lactic acid. Differential physiological and biochemical characteristics of strains AF52-21^T^ and CM04-06^T^ with the closest related species of genus *Faecalibacterium* are listed in the species description and in **Table 1**.

**Table 1.**
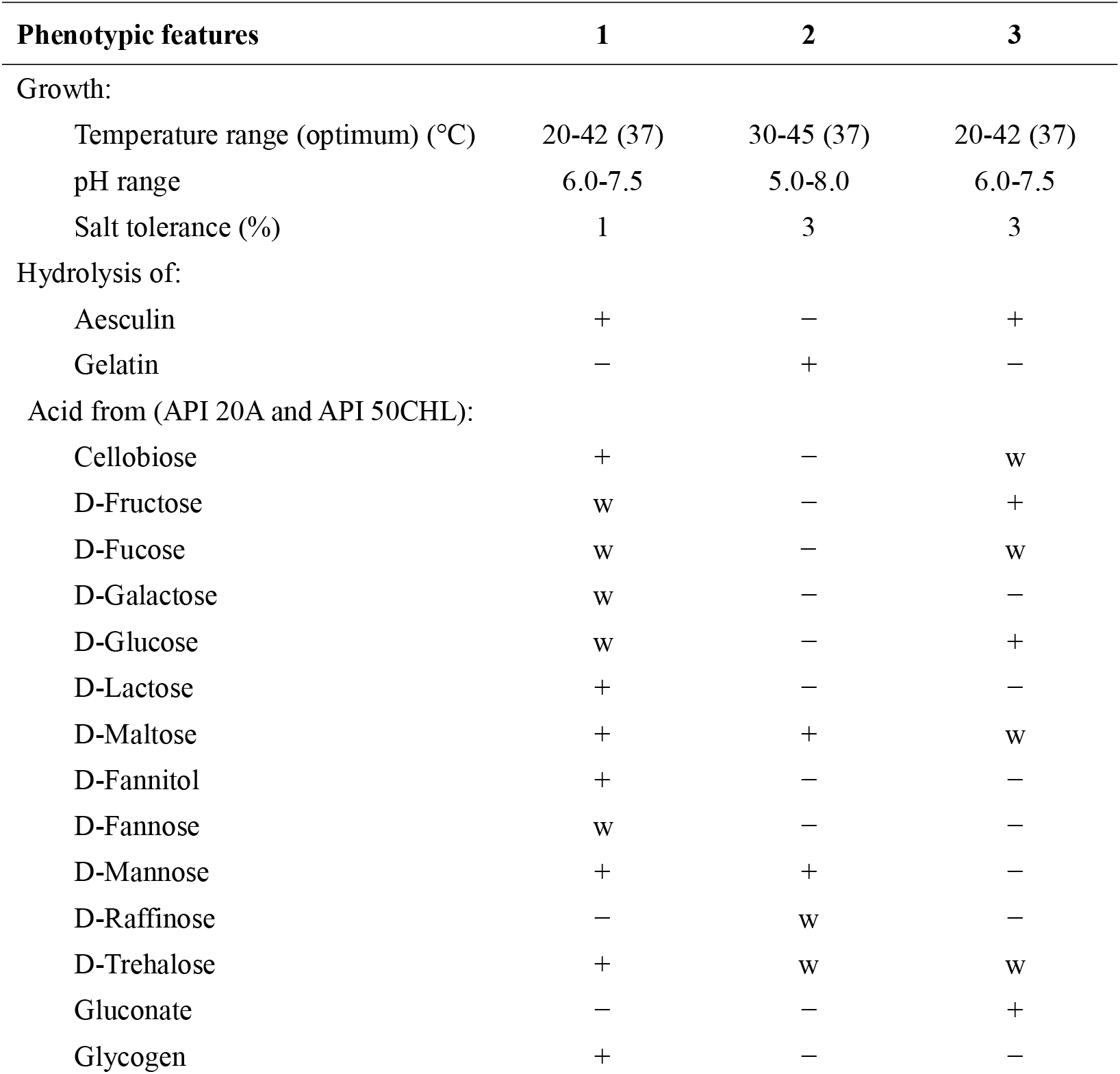
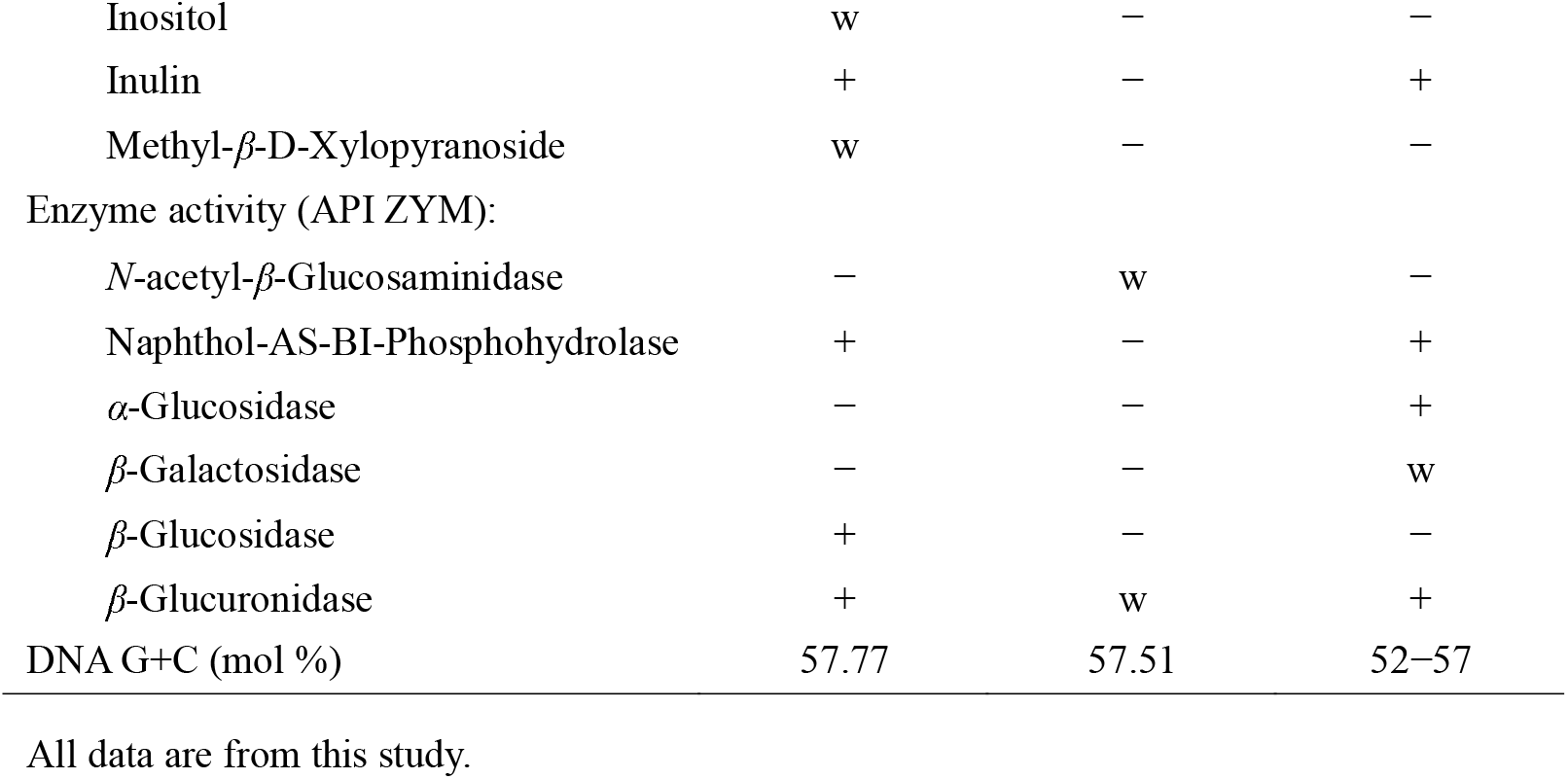
Differential phenotypic characteristics of strains AF52-21^T^, CM04-06^T^, and the related species *F. prausnitzii* ATCC 27768^T^. Strains: 1, *F. butyricigenerans* AF52-21^T^; 2, *F. longum* CM04-06^T^; 3, *F. prausnitzii* ATCC 27768^T^. +, positive; w, weakly positive; –, negative.

| Phenotypic features | 1 | 2 | 3 |
| --- | --- | --- | --- |
| Growth: |  |  |  |
| Temperature range (optimum) (°C) | 20-42 (37) | 30-45 (37) | 20-42 (37) |
| pH range | 6.0-7.5 | 5.0-8.0 | 6.0-7.5 |
| Salt tolerance (%) | 1 | 3 | 3 |
| Hydrolysis of: |  |  |  |
| Aesculin | + | – | + |
| Gelatin | – | + | – |
| Acid from (API 20A and API 50CHL): |  |  |  |
| Cellobiose | + | – | w |
| D-Fructose | w | – | + |
| D-Fucose | w | – | w |
| D-Galactose | w | – | – |
| D-Glucose | w | – | + |
| D-Lactose | + | – | – |
| D-Maltose | + | + | w |
| D-Fannitol | + | – | – |
| D-Fannose | w | – | – |
| D-Mannose | + | + | – |
| D-Raffinose | – | w | – |
| D-Trehalose | + | w | w |
| Gluconate | – | – | + |
| Glycogen | + | – | – |
| Inositol | w | — | — |
| Inulin | + | — | + |
| Methyl-β-D-Xylopyranoside | w | — | — |
| Enzyme activity (API ZYM): |  |  |  |
| *N*-acetyl-β-Glucosaminidase | — | w | — |
| Naphthol-AS-BI-Phosphohydrolase | + | — | + |
| α-Glucosidase | — | — | + |
| β-Galactosidase | — | — | w |
| β-Glucosidase | + | — | — |
| β-Glucuronidase | + | w | + |
| DNA G+C (mol %) | 57.77 | 57.51 | 52–57 |

All data are from this study.

**Figure 1.**
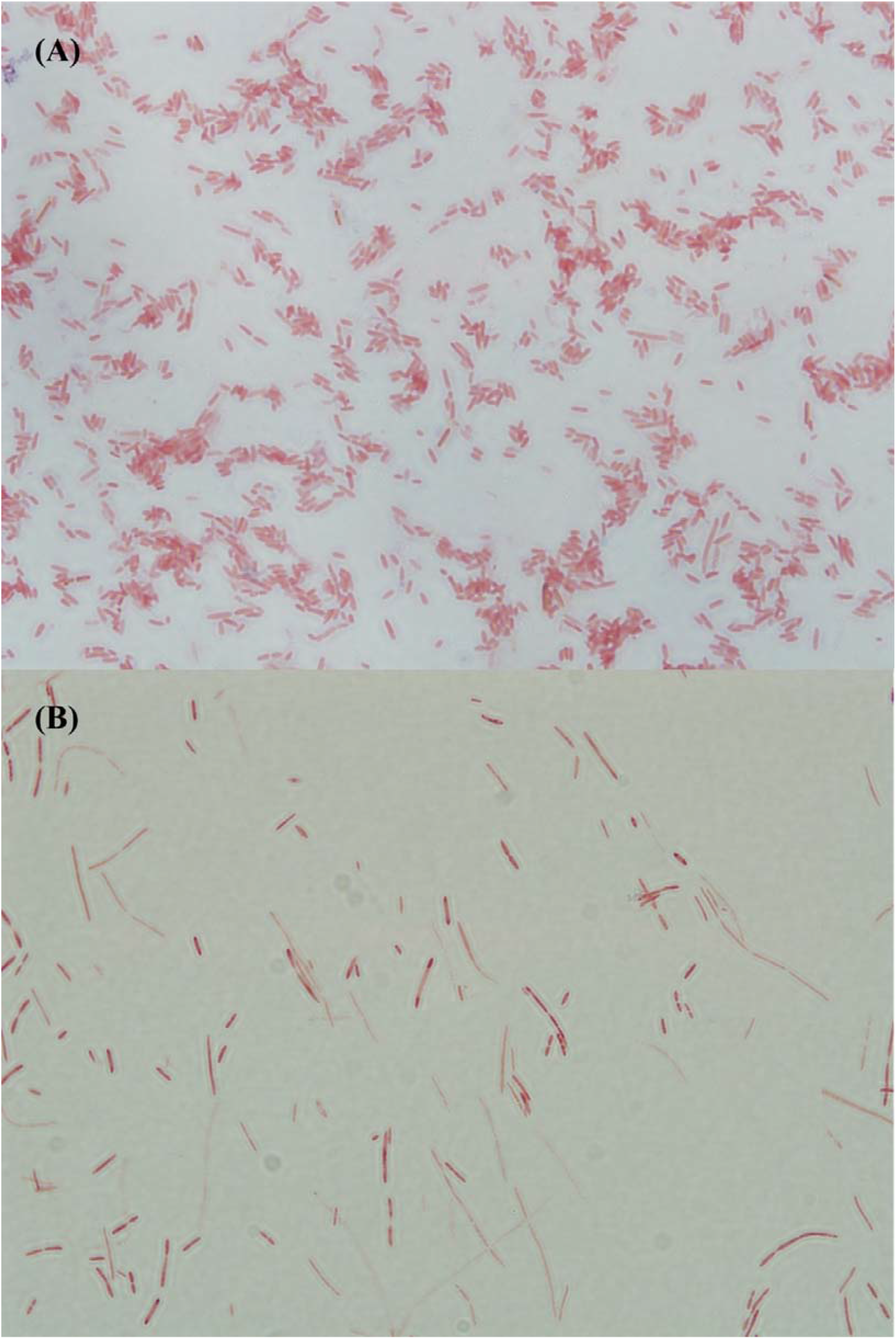
Micrographs of strains AF52-21^T^, CM04-06^T^ after Gram staining. A, AF52-21^T^; B,CM04-06^T^.

The result of cellular fatty acid profiles of strain AF52-21^T^, CM04-06^T^ and related species are shown in **Table 2**. The major components of fatty acids (constituting >5% of the total) present in strain AF52-21^T^ were C_14:0_ (5.9%), C_16:0_ (16.3%), C_18:1_ ω7*c* (8.1), C_18:1_ ω9*c* (39.0%) and iso-C_19:0_(12.9%). The profiles including C_16:0_ (25.5%), C_18:1_ ω7*c* (7.5%), C_18:1_ ω9*c* (32.5%), iso-C_19:0_ (5.9%) and iso-C_17:1_ I/anteiso B (9.7%) were detected as the predominant fatty acids for strain CM04-06^T^. The highest levels of fatty acids, including C_16:0_ and C_18:1_ ω9*c*, were similar, but not identical comparing strain AF52-21^T^, CM04-06^T^ and ATCC 27768^T^. Furthermore, strains AF52-21^T^, CM04-06^T^ and ATCC 27768^T^ could be differentiated by less abundant fatty acids, such as C_18:1_ 2OH, anteiso-C_15:0_, anteiso-C_17:0_, C_13:0_ 3OH/Iso-C_15:1_ I, C_16:1_ ω7*c*/C_16:1_ ω6*c* and antei-C_18:0_ /C_18:2_ ω6, 9*c* (**Table 2**).

**Table 2.** Fatty acid profile of strains AF52-21^T^, CM04-06^T^ and the closest related species *F. prausnitzii* ATCC 27768^T^. Numbers represent percentages of the total fatty acids. –, not detected (<1%).

| Fatty acids composition | *F. butyricigenerans*<br>AF52-21<sup>T</sup> | *F. longum*<br>CM04-06<sup>T</sup> | *F. prausnitzii*<br>ATCC 27768<sup>T</sup> |
| --- | --- | --- | --- |
| C<sub>12:0</sub> | 1.5 | 1.8 | 1.9 |
| C<sub>13:1</sub> | – | – | 1.25 |
| C<sub>14:0</sub> | **5.9** | 4.6 | **11.8** |
| C<sub>16:0</sub> | **16.3** | **25.5** | **21.1** |
| C<sub>17:1</sub> *ω*8*c* | 1.3 | – | 1.1 |
| C<sub>18:1</sub> *ω*7*c* | **8.1** | **7.5** | **5.7** |
| C<sub>18:1</sub> *ω*9*c* | **39.0** | **32.5** | **31.4** |
| C<sub>18:0</sub> | 4.5 | 3.5 | 4.1 |
| C<sub>18:1</sub> 2OH | 2.9 | – | 2.0 |
| Iso-C<sub>19:1</sub> I | 1.2 | 1.1 | 2.1 |
| Iso-C<sub>19:0</sub> | **12.9** | **5.9** | – |
| Anteiso-C<sub>15:0</sub> | – | 2.6 | – |
| Anteiso-C<sub>17:0</sub> | – | 2.1 | – |
| C<sub>13:0</sub> 3OH/ Iso-C<sub>15:1</sub> I | – | – | 2.1 |
| C<sub>16:1</sub> *ω*7*c*/ C<sub>16:1</sub> *ω*6*c* | 1.5 | 1.9 | 4.0 |
| Iso-C<sub>17:1</sub> I/anteiso B | 4.7 | **9.7** | **7.6** |
| Antei-C<sub>18:0</sub>/C<sub>18:2</sub> *ω*6, 9*c* | – | 1.9 | 1.3 |

### Phylogenetic Analysis

The almost complete 16S rRNA gene sequences of strains AF52-21^T^ and CM04-06^T^, comprising 1,382bp and 1,374bp, respectively, were obtained. BLAST analysis of the 16S rRNA gene sequences against the EzBioCloud server showed that the two strains grouped in the genus *Faecalibacterium* within the family *Ruminococcaceae* and were most closely related to *F. prausnitzii* ATCC 27768^T^, which is the sole valid species of the genus *Faecalibacterium*, with similarity values of 97.18% and 96.87%, respectively. *Faecalibacterium hominis* 4P-15^T^, an unrecognized species of the genus *Faecalibacterium*, was also used as a related taxa for 16S rRNA gene analysis. Strains AF52-21^T^ and CM04-06^T^ share 16S rRNA gene sequence similarity of 98.65% and 97.68% with *F. hominis* 4P-15^T^. The 16S rRNA gene sequence similarity between strains AF52-21^T^ and CM04-06^T^ was 98.69% (**Table 3**), both these values were lower than the recommended thresholds (98.7%) for classification of human-associated bacterial isolates at the species level ^32^. Phylogenetic analysis based on the neighbour-joining, maximum-likelihood and minimum-evolution (**Fig. 2, Supplementary Fig. S1** and **Fig. S2**, respectively) confirmed the result of affiliation of the novel isolates with the species of the genus *Faecalibacterium* and revealed that the two isolates formed a distinct lineage with *F. prausnitzii* ATCC 27768^T^.

**Table 3.**
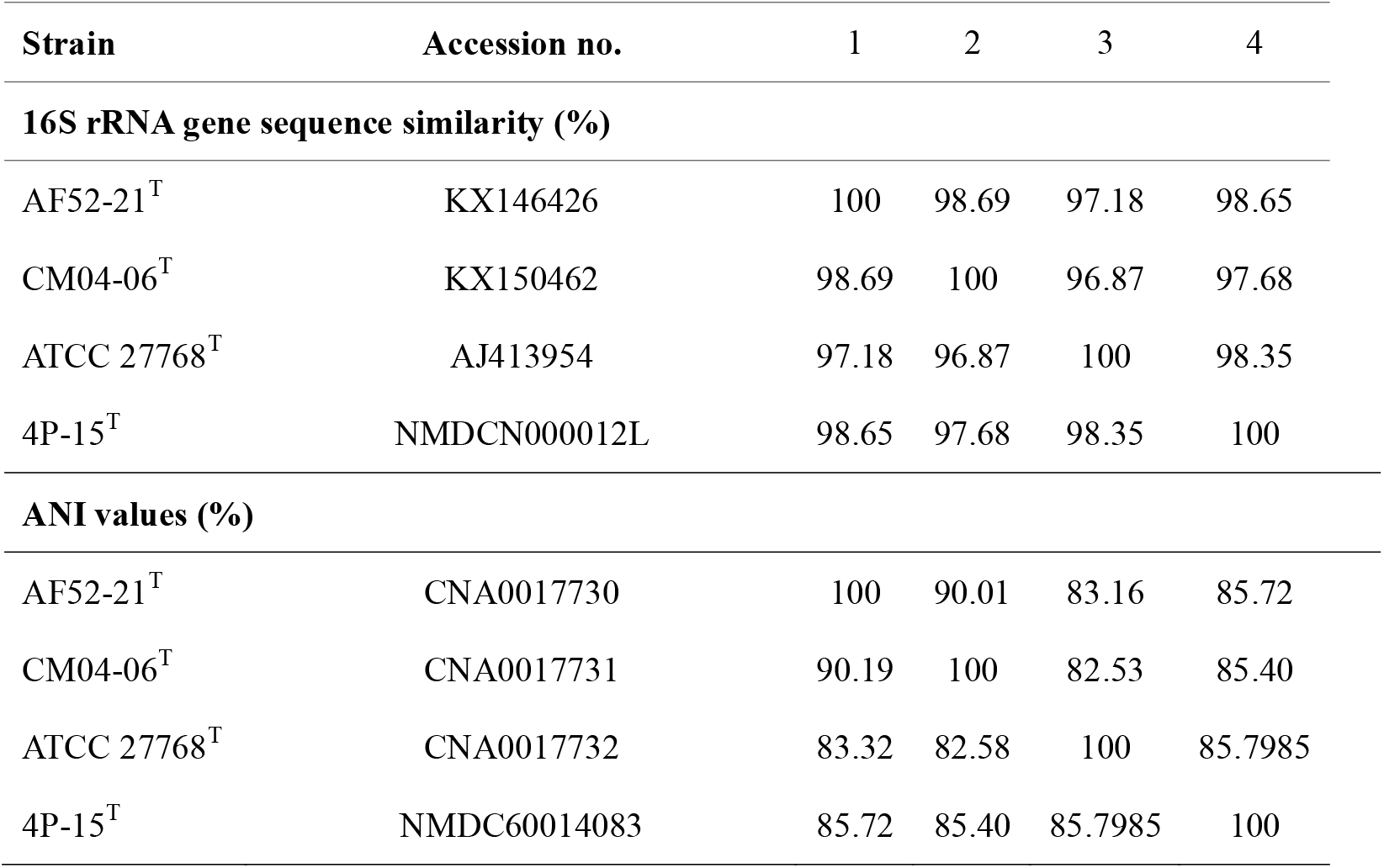
Levels of 16S rRNA gene sequence similarity and ANI values (in percentages) based on BLAST for strains AF52-21^T^, CM04-06^T^ and the phylogenetically related species *F. prausnitzii* ATCC 27768^T^ and the unrecognized species *Faecalibacterium hominis* 4P-15^T^. Taxa: 1, *F. butyricigenerans* AF52-21^T^; 2, *F. longum* CM04-06^T^; 3, *F. prausnitzii* ATCC 27768^T^; 4, *Faecalibacterium hominis* 4P-15^T^.

| Strain | Accession no. | 1 | 2 | 3 | 4 |
| --- | --- | --- | --- | --- | --- |
| **16S rRNA gene sequence similarity (%)** |  |  |  |  |  |
| AF52-21<sup>T</sup> | KX146426 | 100 | 98.69 | 97.18 | 98.65 |
| CM04-06<sup>T</sup> | KX150462 | 98.69 | 100 | 96.87 | 97.68 |
| ATCC 27768<sup>T</sup> | AJ413954 | 97.18 | 96.87 | 100 | 98.35 |
| 4P-15<sup>T</sup> | NMDCN000012L | 98.65 | 97.68 | 98.35 | 100 |
| **ANI values (%)** |  |  |  |  |  |
| AF52-21<sup>T</sup> | CNA0017730 | 100 | 90.01 | 83.16 | 85.72 |
| CM04-06<sup>T</sup> | CNA0017731 | 90.19 | 100 | 82.53 | 85.40 |
| ATCC 27768<sup>T</sup> | CNA0017732 | 83.32 | 82.58 | 100 | 85.7985 |
| 4P-15<sup>T</sup> | NMDC60014083 | 85.72 | 85.40 | 85.7985 | 100 |

**Figure 2.**
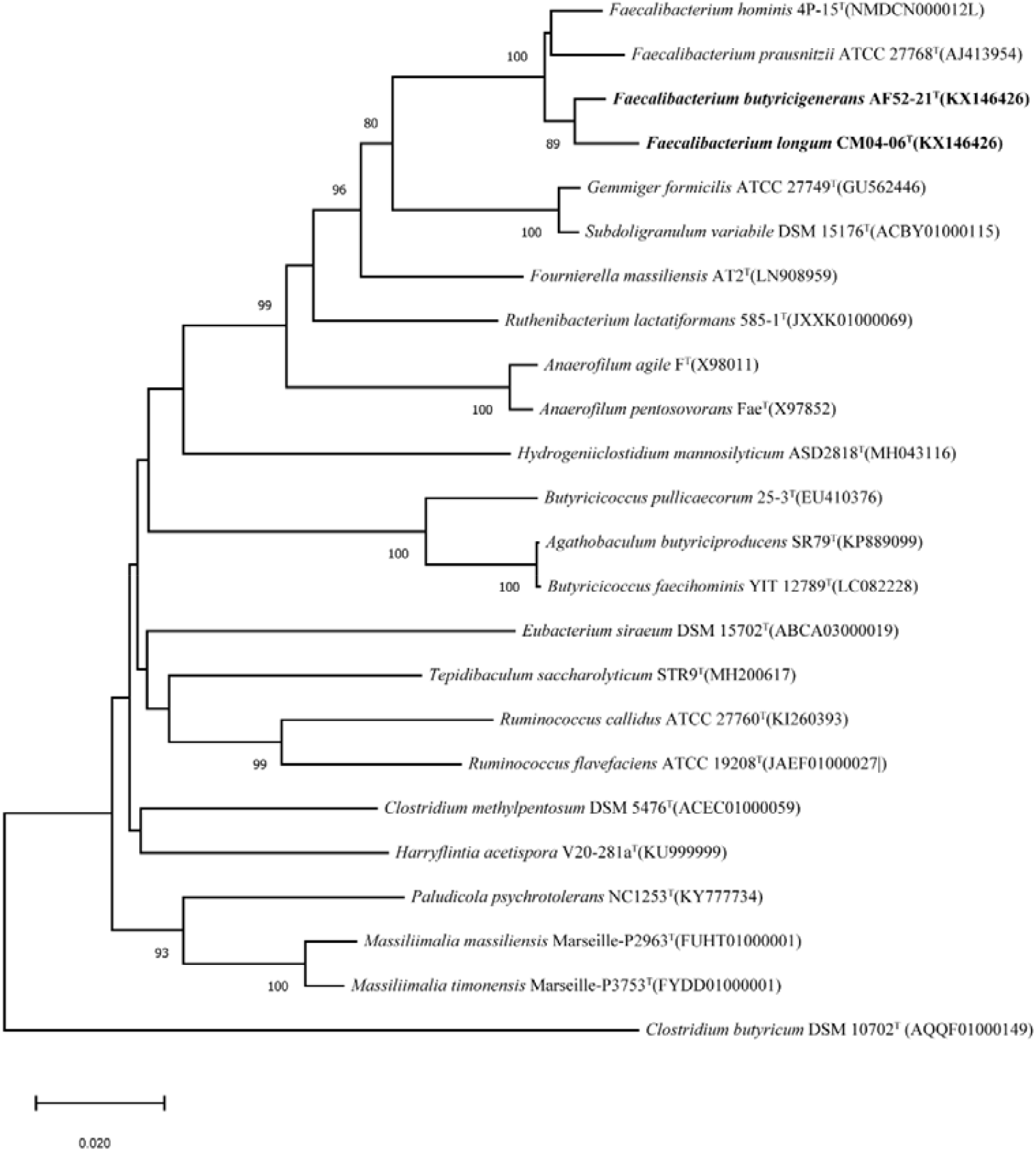
Neighbour-joining phylogenetic tree based on 16S rRNA gene sequences showing the phylogenetic relationships of strains AF52-21^T^, CM04-06^T^ and the representatives of several other related taxa within the family *Ruminococcaceae*. *Clostridium butyricum* DSM 10702^T^ (AQQF01000149) was used as an out-group. Bootstrap values based on 1000 replications higher than 70% are shown at the branching points. Bar, substitutions per nucleotide position.

### Genome Analysis and function annotation

The assembled draft genomes of strains AF52-21^T^ and CM04-06^T^ comprised total lengths of 2,851,918bp and 3,011,178bp with 73 and 47 scaffolds, respectively (**Table 4**). The G+C contents calculated from genome sequences were 57.77% and 57.51%, which were slightly higher than the range reported previously for the genus *Faecalibacterium* (47-57 mol%) ^28^. CheckM analysis of the genomes showed high completeness (>90%) and low contamination (<5%) (**Table 4**), indicating these are high-quality genomes sequences. The genome comparison between strains AF52-21^T^, CM04-06^T^, ATCC 27768^T^ and 4P-15^T^ showed ANI values ranged from 82.53% to 90.19% (**Table 3**), which were significantly below the proposed cutoff value of 95–96% for delineating bacterial species, indicating that strains AF52-21^T^ and CM04-06^T^ represented novel species in the genus *Faecalibacterium*. Circular maps of the two strains AF52-21^T^ and CM04-06^T^ are shown in **Fig. 3** and **Fig. 4**.

**Table 4.**
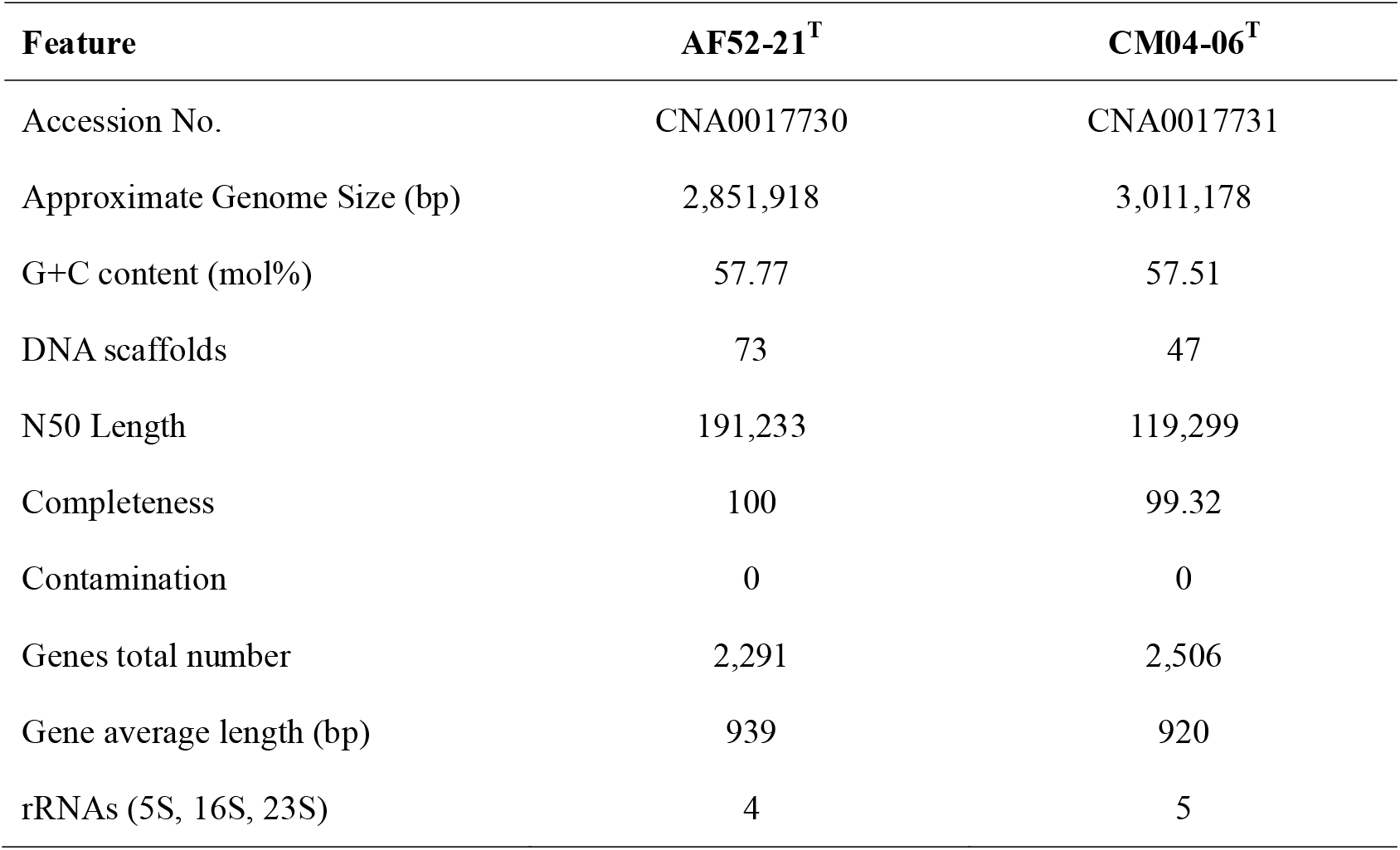
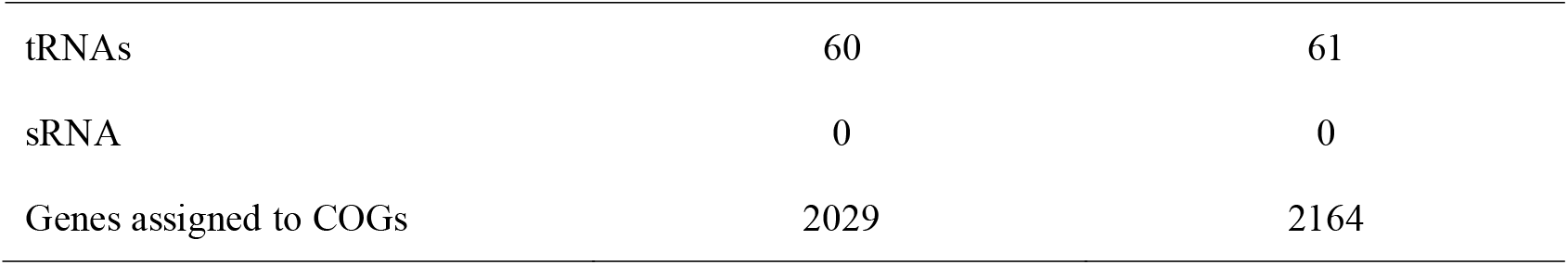
Genome properties of *F. butyricigenerans* AF52-21^T^ and *F. longum* CM04-06^T^.

**Figure 3.**
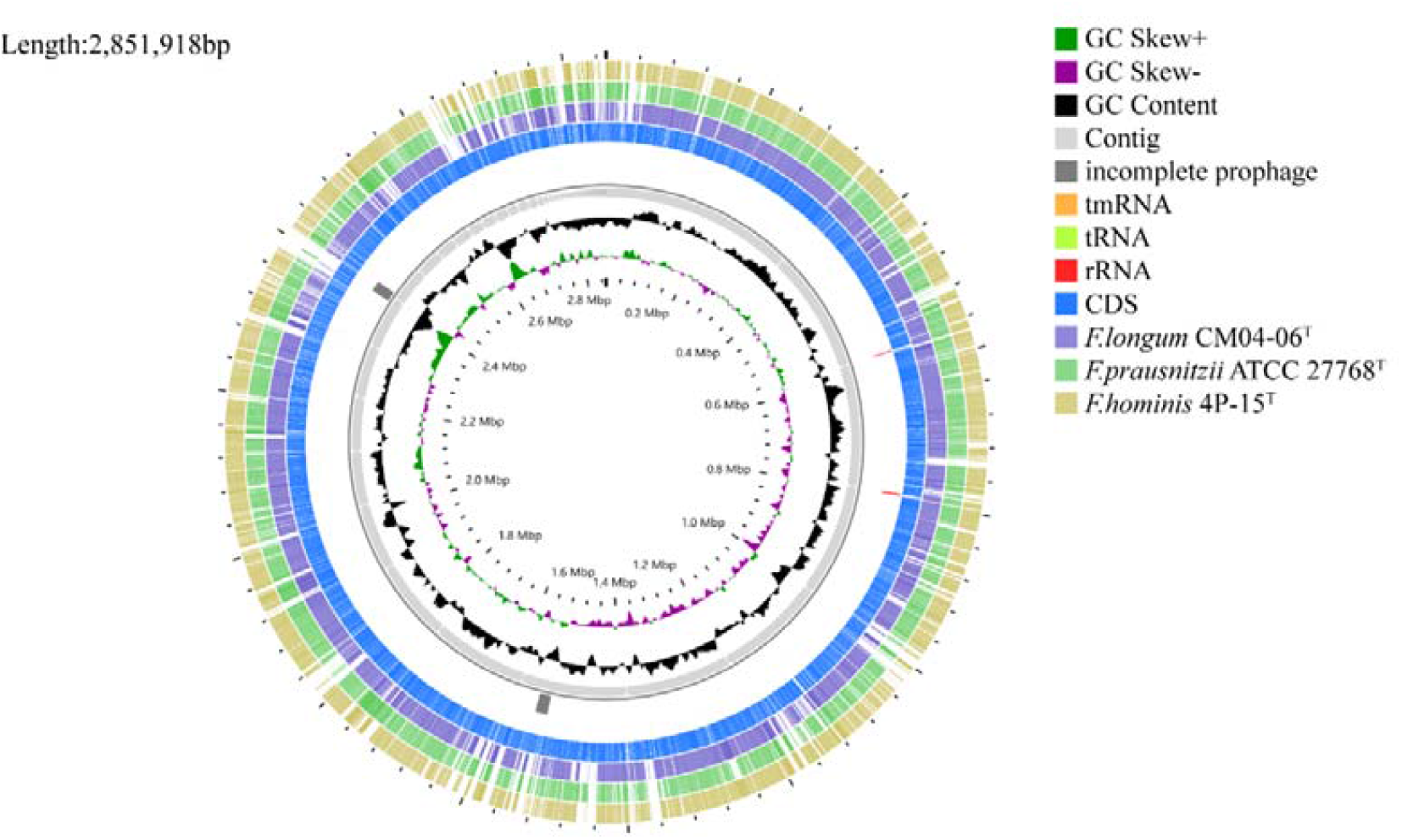
Circular map of AF52-21^T^. Innermost circle, GC skew; circle 2, G+C content; circle 3, contigs; circles 4, predicted prophage remnants; circle 5, tmRNA, tRNA and rRNA genes; circles 6, CDS; circles 7-9, homologous genomic segments from CM04-06 ^T^, *F. prausnitzii* ATCC 27768 ^T^ and *F. hominis* 4P-15^T^.

**Figure 4.**
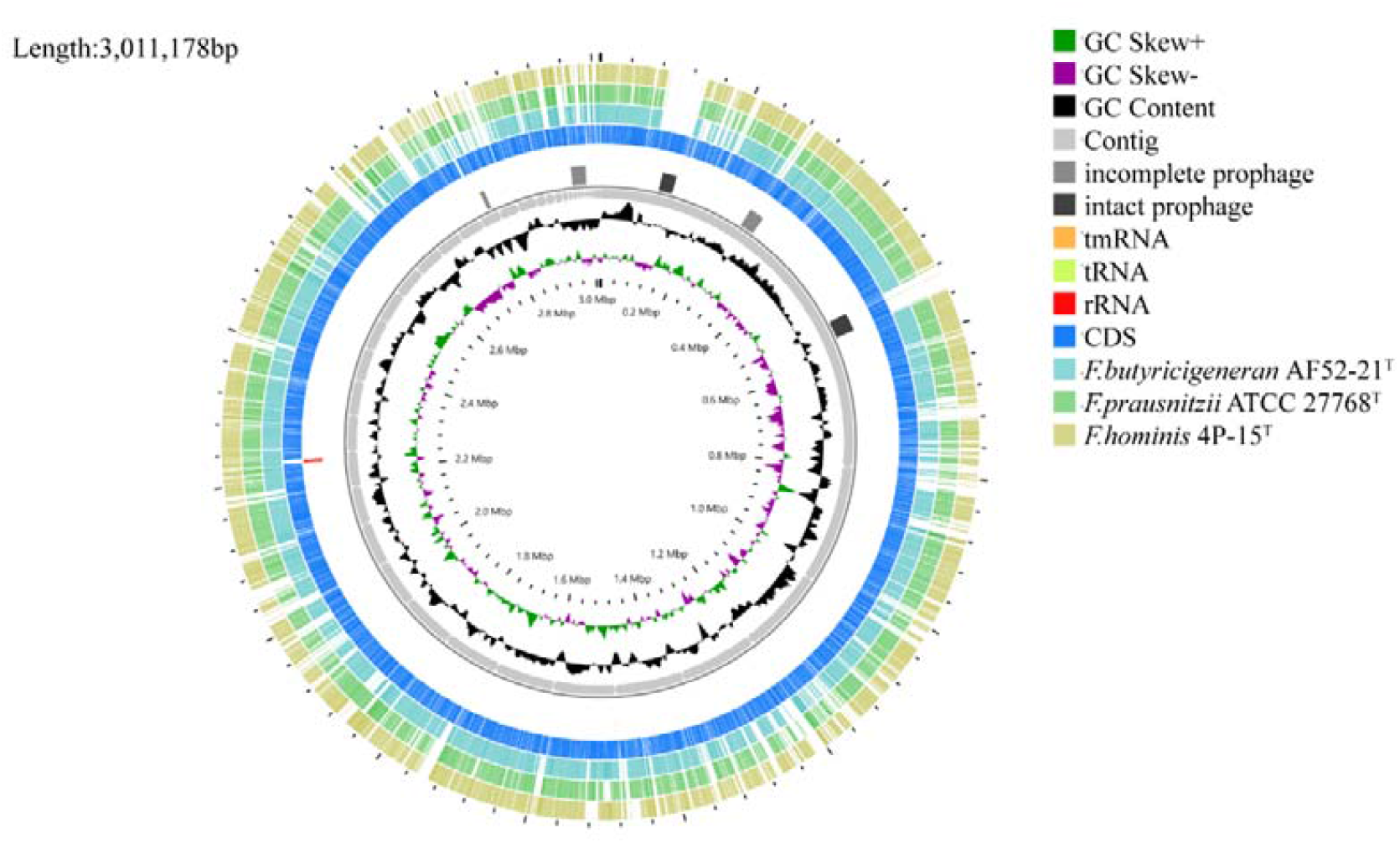
Circular map of CM04-06^T^. Innermost circle, GC skew; circle 2, G+C content; circle 3, contigs; circles 4, predicted prophage remnants; circle 5, tmRNA, tRNA and rRNA genes; circles 6, CDS; circles 7-9, homologous genomic segments from AF52-21 ^T^, *F. prausnitzii* ATCC 27768 ^T^ and *F*.*hominis* 4P-15^T^.

For genome annotation, the distributions of the genes into clusters of orthologous groups (COGs) functional categories are depicted in **Fig. 5** and **Table S1**. Both strain strains AF52-21^T^ and CM04-06^T^ share identical COGs functional categories. The abundant categories comprise amino acid transport and metabolism (E), carbohydrate transport and metabolism (G), cell wall/membrane/envelope biogenesis (M), energy production and conversion (C), general function prediction only (R), replication, recombination and repair (L), signal transduction mechanisms (T), transcription (K) and translation, ribosomal structure and biogenesis (J). Annotated genes associated with synthesis of diaminopimelic acid, teichoic and lipoteichoic acids and lipopolysaccharides, and metabolism of polar lipids and polyamines by RAST annotation, comparing strains AF52-21^T^, CM04-06^T^ with ATCC 27768^T^, are shown in **Table 5** and **Table S2**.

**Table 5.**
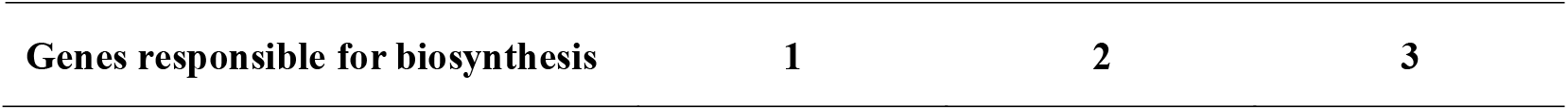
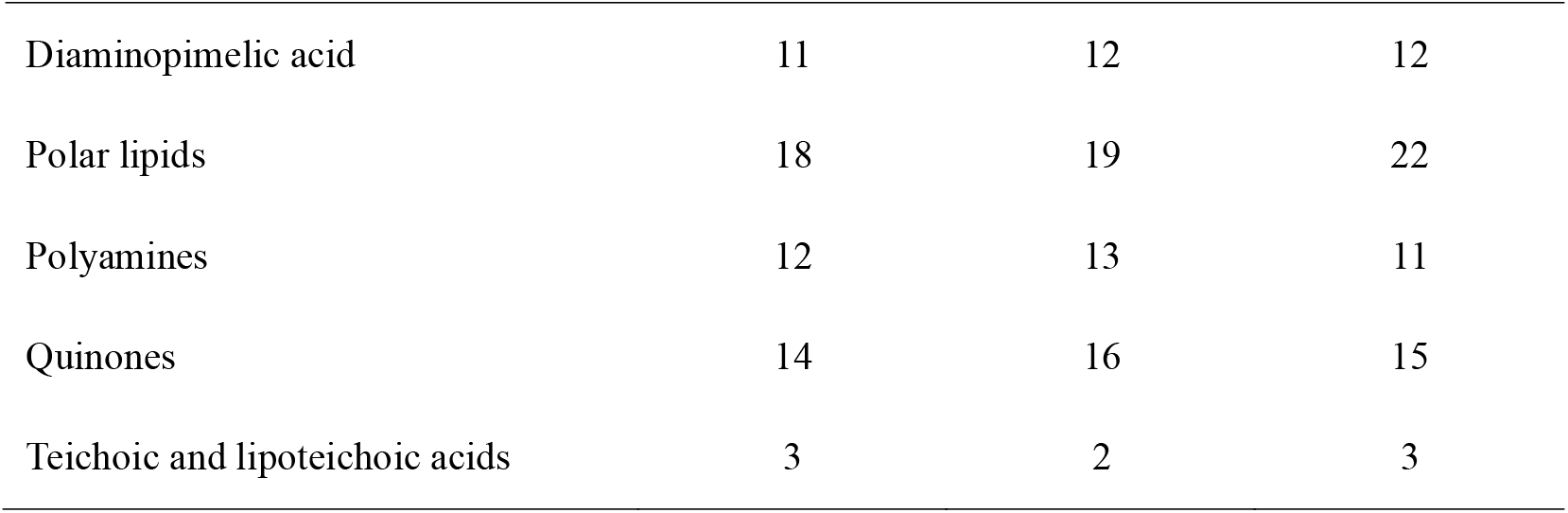
Number of genes associated with biosynthetic pathway from whole genome sequences of strain *F. butyricigenerans* AF52-21^T^ and *F. longum* CM04-06^T^ and *F. prausnitzii* ATCC 27768^T^ identified by RAST. Taxa: 1, AF52-21^T^; 2, CM04-06^T^; 3, *F. prausnitzii* ATCC 27768^T^. Data are for type strains. Numbers of genes identified for lipopolysaccharides and mycolic acids were zero for all taxa studied.

**Figure 5.**
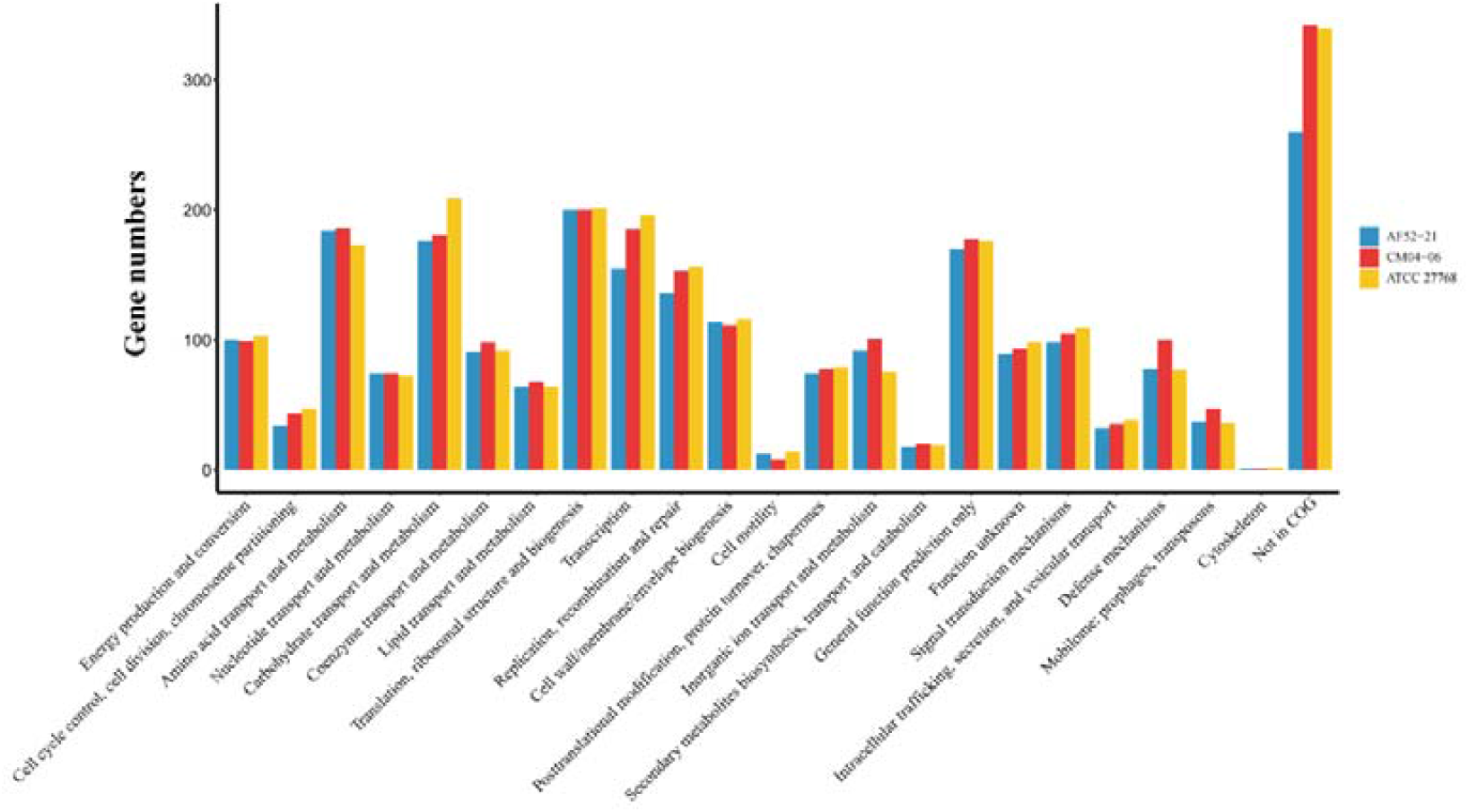
Comparison of COG functional categories of strains AF52-21^T^, CM04-06^T^ and the closest related species *F. prausnitzii* ATCC 27768^T^.

The annotation showed that AF52-21^T^, CM04-06^T^, and ATCC 27768^T^ contained a complete acetyl-CoA to butyrate synthesis pathway, but possessed butyryl-CoA:acetate CoA-transferase activity only in the final step (**Fig. 6**), as discussed previously ^33,34^. Prophages were identified using the PHAST software, and the results are shown in **Fig. 7**. Two incomplete phage sequences were detected in the AF52-21^T^ genome, one of which encodes the Phd_YefM protein, an antitoxin component. Three incomplete phage sequences and two intact prophages were detected in the CM04-06^T^ genome, encoding the Phd_YefM protein, relaxase/mobilisation nuclease domain, bacterial mobilisation protein (MobC) /ribbon-helix-helix protein, helix-turn-helix, and predicted transcriptional regulators. Moreover, the antibiotic resistance analysis indicated that AF52-21^T^ contained macrolide antibiotic, lincosamide antibiotic, and streptogramin antibiotic genes, while CM04-06^T^ and ATCC 27768^T^ contained aminoglycoside antibiotic genes (**Fig. 8**).

**Figure 6.**
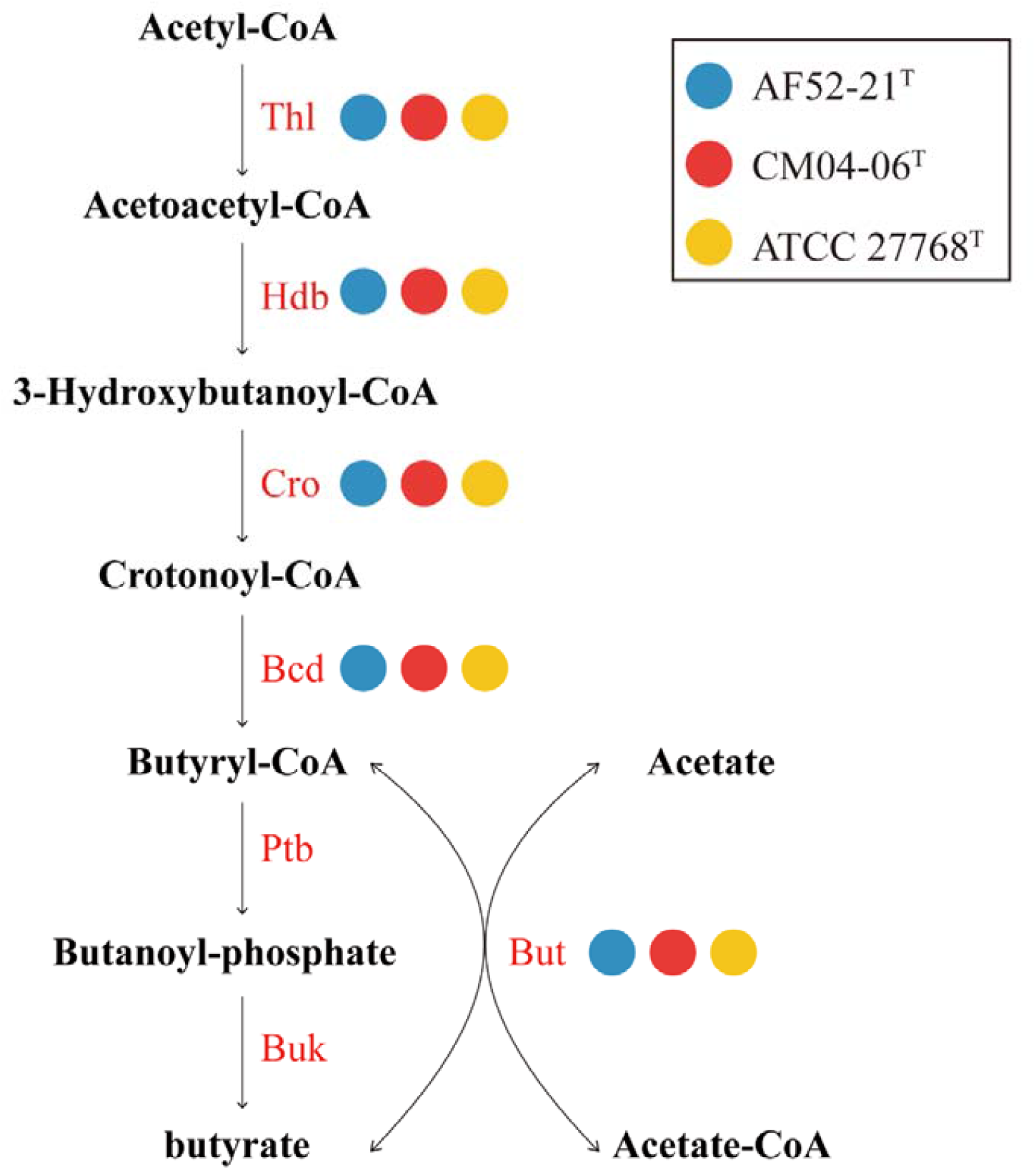
The synthesis pathways from Acetyl-CoA to Butyrate. Strains AF52-21^T^, CM04-06^T^ and ATCC 27768^T^ were annotated as blue, red, and yellow. Thl, thiolase; Hdb, *β*-hydroxybutyryl-CoA dehydrogenase; Cro, crotonase; Bcd, butyryl-CoA dehydrogenase; But, butyryl-CoA:acetate CoA transferase; Ptb, phosphate butyryltransferase; Buk, butyrate kinase.

**Figure 7.**
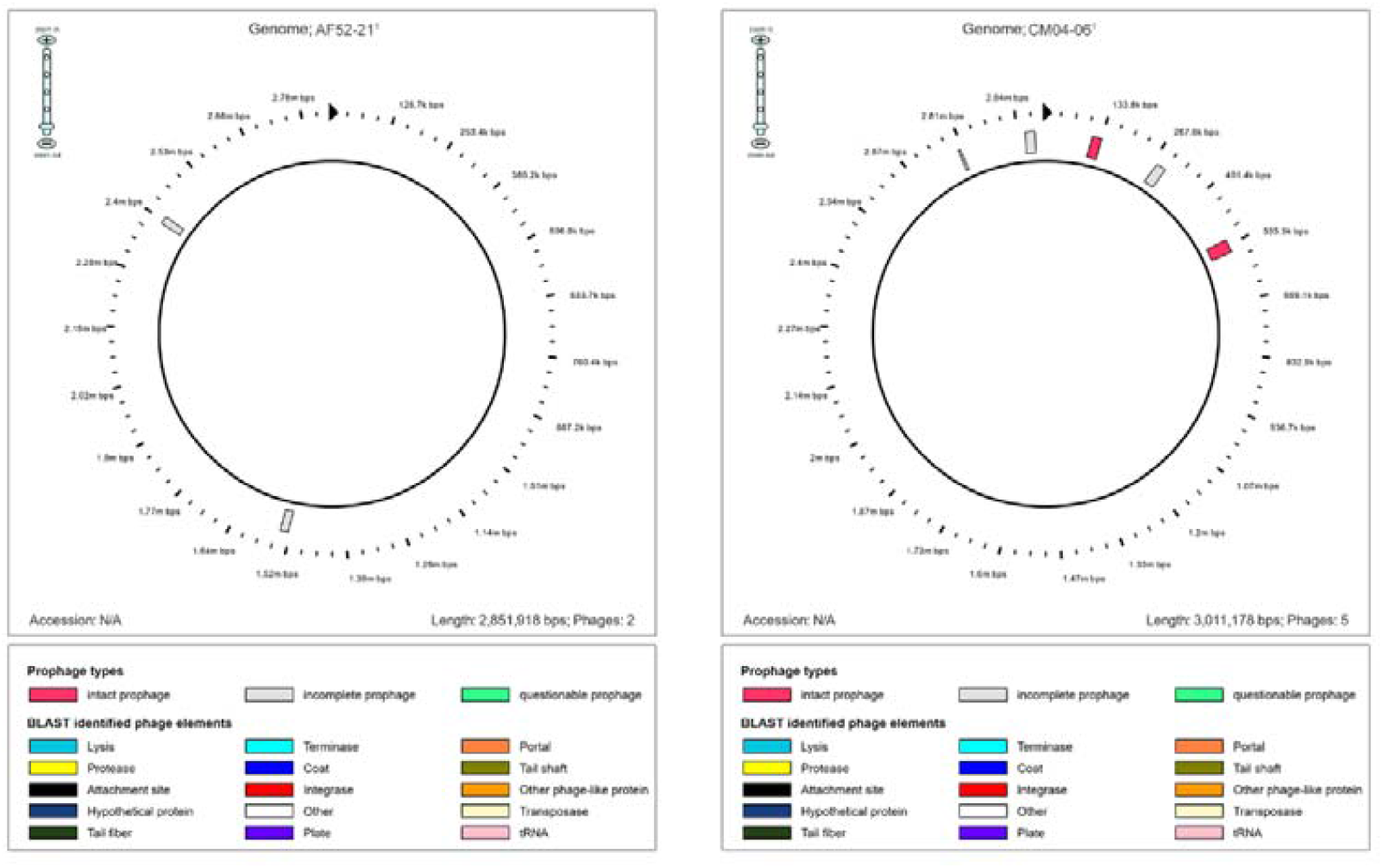
Distribution of prophage in strains AF52-21T and CM04-06T.

**Figure 8.**
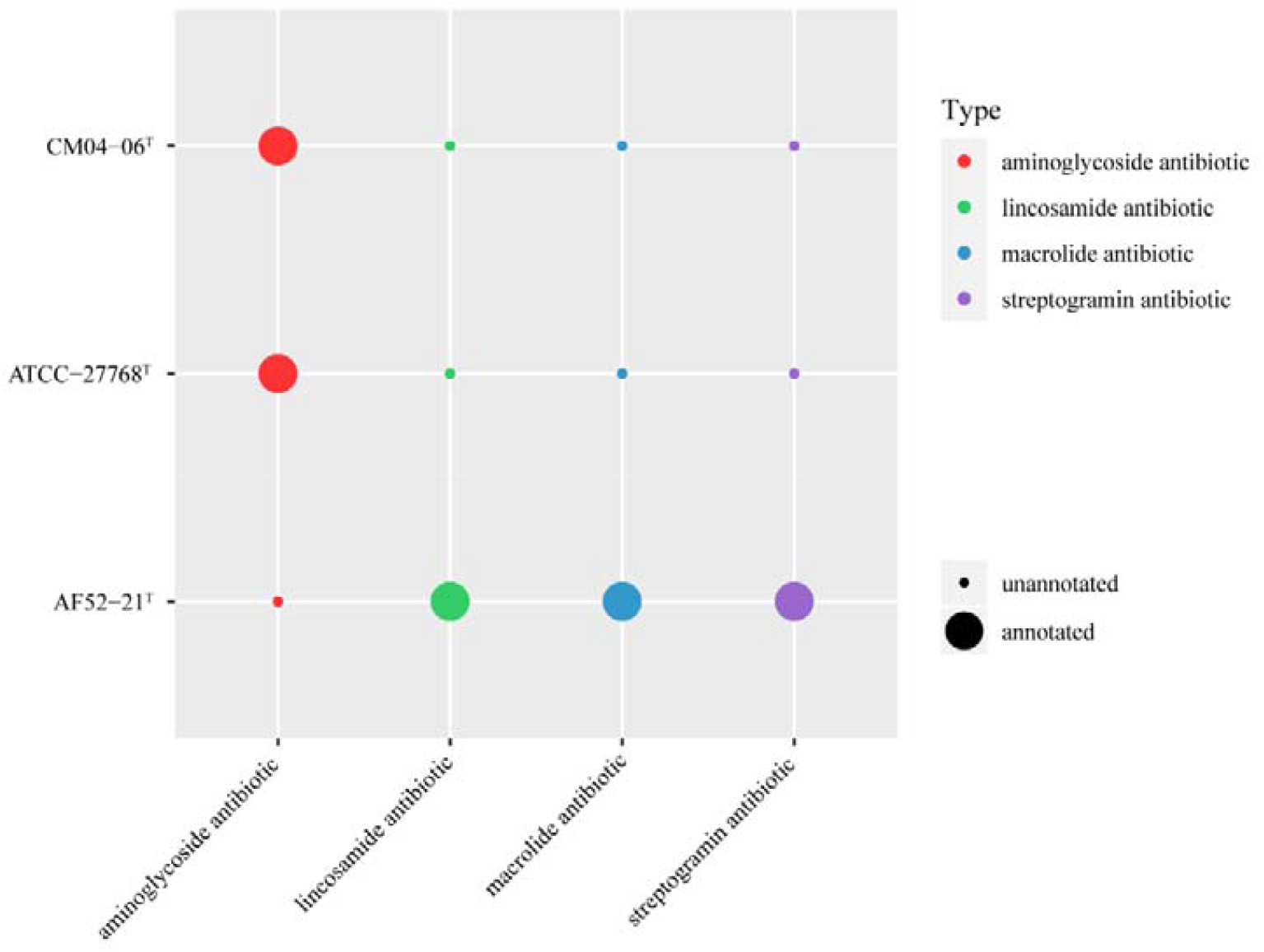
Comparison of resistance genes in strains AF52-21^T^, CM04-06^T^, and *F. prausnitzii* ATCC 27768^T^.

## Discussion

The 16S rRNA genes phylogenetic, physiological results and genome description showed that the two new isolates AF52-21^T^ and CM04-06^T^ represent two novel species. The ANI values between AF52-21^T^, CM04-06^T^ and the closest related species ATCC 27768^T^ were 82.54% and 90.09%, respectively, which is in support of a new species delineation. The result of biochemical and genomic functional analyses showed that both AF52-21^T^ and CM04-06^T^ are butyric acid-producing bacteria.

Most strains in the genus *Faecalibacterium* exhibit a common ability to produce butyric acid, peptides and other anti-inflammatory substances, which have immunomodulatory effects^26,27,35^. Some studies have confirmed that the decreased abundance of this genus is related to the occurrence and development of inflammatory bowel diseases^36-38^. Accordingly, *Faecalibacterium* is receiving much attention as one of the candidate next-generation probiotics (NGPs), which can be used for disease treatment^39,40^.

Previous studies based on comparative genomics from isolates suggested the wide diversity of this genus, with the presence of at least two phylotypes in *F. prausnitzii*^29^. A recent study analysing the *Faecalibacterium*-like MAGs, proposed that *Faecalibacterium* from the human gut can be divided into 12 clades^40^. These studies have expanded the diversity of *Faecalibacterium* and proposed that different phylotypes have different functions, which results in different contributions to health or diseases.

Moreover, as a candidate taxa for the NGPs, the *Faecalibacterium* isolates can be used for in-vitro functional verification and animal model experiments to further explore its probiotic functions, and ultimately expected to be used in clinical disease intervention.

### Description of Faecalibacterium butyricigenerans sp. nov

*Faecalibacterium butyricigenerans* (bu.ty.ri.ci.ge′ne.rans. N.L. n. *acidum butyricum* butyric acid; L. part. adj. *generans*, producing; N.L. adj. *butyricigenerans*, butyric acid-producing; referring to its production of butyric acid)

Cells of strain AF52-21^T^ are Gram-stain-negative, non-motile, non-spore-forming and rod-shaped. Strictly anaerobic and catalase negative. Colonies on PYG agar are round, creamy white to yellowish, convex and opaque with entire margins and colony size is approximately 1.0-2.0 mm in diameter after incubation at 37°C for 2 days. Cells are able to grow at 20-42°C with optimum temperature at 37°C. The pH range for growth is 6.0-7.5 (optimum at 7.0-7.5). Growth occurs at NaCl concentrations 0-1%. The strain is positive for the assimilation of cellobiose, D-lactose, D-maltose, D-mannitol, D-mannose, D-trehalose, glycogen, inulin, D-fructose, D-fucose, D-galactose, D-glucose, inositol and methyl-*β*-D-xylopyranoside, but negative for amygdalin, arbutin, D-adonitol, D-arabinose, D-arabitol, D-lyxose, D-melezitose, D-melibiose, D-raffinose, D-ribose, D-sorbitol, D-tagatose, D-turanose, dulcitol, D-xylose, erythritol, gentiobiose, gluconate, glycerol, L-arabinose, L-arabitol, L-fucose, L-rhamnose, L-sorbose, L-xylose, methyl-D-glucopyranoside, methyl-*α*-D-mannopyranoside, *N*-acetyl-glucosamine, salicin, starch, sucrose, xylitol, 2-ketogluconate and 5-ketogluconate. In enzymatic activity tests, strain AF52-21^T^ is positive for naphthol-AS-BI-phosphohydrolase, *β*-glucuronidase and *β*-glucosidase, and negative for alkaline phosphatase, esterase (C4), esterase lipase (C8), lipase (C14), leucine arylamidase, valine arylamidase, cystine arylamidase, trypsin, *α*-chymotrypsin, acid phosphatase, naphthol-AS-BI-phosphohydrolase, *α*-galactosidase, *β*-galactosidase, *α*-glucosidase, *α*-mannosidase and *β*-fucosidase. Indole is not produced. Positive for hydrolysis of esculin and negative for gelatin. Formic acid, acetic acid, butyric acid and lactic acid are the fermentation products. The major fatty acids are C_14:0_, C_16:0_, C_18:1_ ω7*c*, C_18:1_ ω9*c* and iso-C_19:0_.

In the result of RAST annotation, 11 genes/proteins are associated with biosynthesis of DAP, including 4-hydroxy-tetrahydrodipicolinate reductase (EC 1.17.1.8), 4-hydroxy-tetrahydrodipicolinate synthase (EC 4.3.3.7), aspartate-semialdehyde dehydrogenase (EC 1.2.1.11), aspartokinase (EC 2.7.2.4) (2 copies), diaminopimelate decarboxylase (EC 4.1.1.20), diaminopimelate epimerase (EC 5.1.1.7), L, L-diaminopimelate aminotransferase (EC 2.6.1.83), *N*-acetyl-L, L-diaminopimelate deacetylase (EC 3.5.1.47), UDP-*N*-acetylmuramoylalanyl-D-glutamate-2, 6-diaminopimelate ligase (EC 6.3.2.13) and UDP-*N*-acetylmuramoylalanyl-D-glutamyl-2, 6-diaminopimelate-D-alanyl-D-alanine ligase (EC 6.3.2.10). 18 genes/proteins are associated with biosynthesis of polar lipids, including 1-acyl-sn-glycerol-3-phosphate acyltransferase (EC 2.3.1.51) (2 copies), ABC-type multidrug/protein/lipid transport system, ATPase component, acyl carrier protein (3 copies), acyl-phosphate:glycerol-3-phosphate O-acyltransferase PlsY, cardiolipin synthetase (EC 2.7.8.-) (3 copies), CDP-diacylglycerol--glycerol-3-phosphate 3-phosphatidyltransferase (EC 2.7.8.5), dihydroxyacetone kinase family protein, glycerol kinase (EC 2.7.1.30), glycerol-3-phosphate dehydrogenase [NAD(P)^+^] (EC 1.1.1.94), phosphate:acyl-ACP acyltransferase PlsX, phosphatidate cytidylyltransferase (EC 2.7.7.41) and phosphatidylglycerophosphatase B (EC 3.1.3.27) (2 copies). 12 genes/proteins are associated with biosynthesis of polyamines, including 5’-methylthioadenosine nucleosidase (EC 3.2.2.16), S-adenosylhomocysteine nucleosidase (EC 3.2.2.9), ABC transporter, periplasmic spermidine putrescine-binding protein PotD (TC 3.A.1.11.1), agmatinase (EC 3.5.3.11), arginine decarboxylase (EC 4.1.1.19), arginine/ornithine antiporter ArcD (2 copies), carboxynorspermidine decarboxylase, putative (EC 4.1.1.-), carboxynorspermidine dehydrogenase, putative (EC 1.1.1.-), putrescine transport ATP-binding protein PotA (TC 3.A.1.11.1), spermidine putrescine ABC transporter permease component PotB (TC 3.A.1.11.1), spermidine putrescine ABC transporter permease component potC (TC..3.A.1.11.1) and spermidine synthase (EC 2.5.1.16). 3 genes/proteins are associated with biosynthesis of teichoic and lipoteichoic acids, including cell wall teichoic acid glycosylation protein gtcA, teichoic acid export ATP-binding protein TagH (EC 3.6.3.40) and membrane protein involved in the export of O-antigen, teichoic acid lipoteichoic acids. 14 genes/proteins are associated with biosynthesis of quinones, including 2-heptaprenyl-1,4-naphthoquinone methyltransferase (EC 2.1.1.163), electron transport complex protein RnfA (2 copies), electron transport complex protein RnfB, electron transport complex protein RnfC, electron transport complex protein RnfD (2 copies), electron transport complex protein RnfE (2 copies), electron transport complex protein RnfG, F420H2:quinone oxidoreductase, heptaprenyl diphosphate synthase component I (EC 2.5.1.30), microsomal dipeptidase (EC 3.4.13.19) and undecaprenyl diphosphate synthase (EC 2.5.1.31). There are no genes responsible for biosynthesis of lipopolysaccharides or mycolic acids. Additional annotations showed that the AF52-21^T^ genome contains a complete butyrate synthesis pathway, two prophage remnants, and three antibiotic genes.

The type strain, AF52-21^T^ (=CGMCC 1.5206^T^ = DSM 103434^T^), was isolated from human gut. The G+C content of the genomic DNA is 57.77 mol% as calculated from whole genome sequencing.

### Description of Faecalibacterium longum sp. nov

#### *Faecalibacterium longum* (lon′gum. L. neut. adj. *longum* long, the shape of the cells)

Cells are Gram-stain-negative, non-motile, non-spore forming, long rod in shape. Strictly anaerobic. Catalase and urease are negative. Colonies are round, yellowish, slightly convex, opaque with entire margins with 2.0 mm in diameter on PYG agar for incubation at 37°C for 48 h under anaerobic condition. The strain showed growth at 30-45°C (optimum temperature is 37°C). Growth is observed at pH 5.0-8.0 (optimum pH is 7.0-7.5). NaCl is tolerated with concentrations up to 3%. Acid is produced from D-maltose, D-mannose, D-raffinose, D-trehalose and salicin, but not from amygdalin, arbutin, cellobiose, D-adonitol, D-arabinose, D-arabitol, D-cellobiose, D-fructose, D-fucose, D-galactose, D-glucose, D-lactose, D-lyxose, D-maltose, D-mannitol, D-mannose, D-melezitose, D-melibiose, D-raffinose, D-ribose, D-sorbitol, D-sucrose, D-tagatose, D-turanose, dulcitol, D-xylose, erythritol, gentiobiose, gluconate, glycerol, glycogen, inositol, inulin, L-arabinose, L-arabitol, L-fucose, L-rhamnose, L-sorbose, L-xylose, methyl-D-glucopyranoside, methyl-*α*-D-mannopyranoside, methyl-*β*-D-xylopyranoside, *N*-acetyl-glucosamine, salicin, starch, sucrose, xylitol, 2-ketogluconate and 5-ketogluconate. In the API ZYM strip, strain showed weakly positive enzyme activities for *β*-glucuronidase and *N*-acetyl-*β*-glucosaminidase, but negative for alkaline phosphatase, esterase (C4), esterase lipase (C8), lipase (C14), leucine arylamidase, valine arylamidase, cystine arylamidase, trypsin, *α*-chymotrypsin, acid phosphatase, naphthol-AS-BI-phosphohydrolase, *α*-galactosidase, *β*-galactosidase, *α*-glucosidase, *β*-glucosidase, *α*-mannosidase and *β*-fucosidase. Indole is not produced. Gelatin is hydrolysed, but aesculin is not. Major end products are acetic acid, formic acid, butyric acid and lactic acid. The major fatty acids (constituting >5% of the total) are C_16:0_, C_18:1_ ω7*c*, C_18:1_ ω9*c*, iso-C_19:0_ and iso-C_17:1_ I/anteiso B.

In the result of RAST annotation, 12 genes/proteins are associated with biosynthesis of DAP, including 4-hydroxy-tetrahydrodipicolinate reductase (EC 1.17.1.8), 4-hydroxy-tetrahydrodipicolinate synthase (EC 4.3.3.7) (2 copies), aspartate-semialdehyde dehydrogenase (EC 1.2.1.11), aspartokinase (EC 2.7.2.4) (2 copies), diaminopimelate decarboxylase (EC 4.1.1.20), diaminopimelate epimerase (EC 5.1.1.7), L, L-diaminopimelate aminotransferase (EC 2.6.1.83), *N*-acetyl-L, L-diaminopimelate deacetylase (EC 3.5.1.47), UDP-*N*-acetylmuramoylalanyl-D-glutamate-2, 6-diaminopimelate ligase (EC 6.3.2.13) and UDP-*N*-acetylmuramoylalanyl-D-glutamyl-2, 6-diaminopimelate-D-alanyl-D-alanine ligase (EC 6.3.2.10). 19 genes/proteins are associated with biosynthesis of polar lipids, including 1-acyl-sn-glycerol-3-phosphate acyltransferase (EC 2.3.1.51) (2 copies), ABC-type multidrug/protein/lipid transport system, ATPase component, acyl carrier protein (3 copies), acyl-phosphate:glycerol-3-phosphate O-acyltransferase PlsY, cardiolipin synthetase (EC 2.7.8.-) (2 copies), CDP-diacylglycerol--glycerol-3-phosphate 3-phosphatidyltransferase (EC 2.7.8.5), dihydroxyacetone kinase family protein, glycerate kinase (EC 2.7.1.31), glycerol kinase (EC 2.7.1.30), glycerol-3-phosphate dehydrogenase [NAD(P)^+^] (EC 1.1.1.94), octaprenyl diphosphate synthase (EC 2.5.1.90) / gimethylallyltransferase (EC 2.5.1.1) / (2E,6E)-farnesyl diphosphate synthase (EC 2.5.1.10) / geranylgeranyl pyrophosphate synthetase (EC 2.5.1.29), phosphate:acyl-ACP acyltransferase PlsX, phosphatidate cytidylyltransferase (EC 2.7.7.41) and phosphatidylglycerophosphatase B (EC 3.1.3.27) (2 copies). 13 genes/proteins are associated with biosynthesis of polyamines, including 5’-methylthioadenosine nucleosidase (EC 3.2.2.16) @ S-adenosylhomocysteine nucleosidase (EC 3.2.2.9), ABC transporter, periplasmic spermidine putrescine-binding protein PotD (TC 3.A.1.11.1), agmatinase (EC 3.5.3.11), arginine decarboxylase (EC 4.1.1.19), arginine/ornithine antiporter ArcD (3 copies), carboxynorspermidine decarboxylase, putative (EC 4.1.1.-), carboxynorspermidine dehydrogenase, putative (EC 1.1.1.-), putrescine transport ATP-binding protein PotA (TC 3.A.1.11.1), spermidine putrescine ABC transporter permease component PotB (TC 3.A.1.11.1), spermidine putrescine ABC transporter permease component potC (TC..3.A.1.11.1) and spermidine synthase (EC 2.5.1.16). 2 genes/proteins are associated with biosynthesis of teichoic and lipoteichoic acids, including cell wall teichoic acid glycosylation protein gtcA and teichoic acid export ATP-binding protein TagH (EC 3.6.3.40). 16 genes/proteins are associated with biosynthesis of quinones, including 2-heptaprenyl-1,4-naphthoquinone methyltransferase (EC 2.1.1.163), electron transport complex protein RnfA (2 copies), electron transport complex protein RnfB, electron transport complex protein RnfC, electron transport complex protein RnfD (2 copies), electron transport complex protein RnfE (2 copies), electron transport complex protein RnfG, heptaprenyl diphosphate synthase component I (EC 2.5.1.30), microsomal dipeptidase (EC 3.4.13.19), octaprenyl diphosphate synthase (EC 2.5.1.90) / dimethylallyltransferase (EC 2.5.1.1) / (2E,6E)-farnesyl diphosphate synthase (EC 2.5.1.10) / geranylgeranyl pyrophosphate synthetase (EC 2.5.1.29) ubiquinone/menaquinone biosynthesis methyltransferase UbiE (EC 2.1.1.-) @ 2-heptaprenyl-1,4-naphthoquinone methyltransferase MenG (EC 2.1.1.163) (2 copies) and undecaprenyl diphosphate synthase (EC 2.5.1.31). There are no genes responsible for biosynthesis of lipopolysaccharides or mycolic acids. Additional annotations showed that the CM04-06^T^ genome contains a complete butyrate synthesis pathway, three phage remnants, two intact prophages, and aminoglycoside antibiotic genes.

The type strain, CM04-06^T^ (=CGMCC 1.5208^T^ = DSM 103432^T^), was isolated from human gut. The G+C content of the genomic DNA is 57.51 mol% as calculated from whole genome sequencing.

## Materials and Methods

### Origin of bacterial strains

Faeces samples were collected from two healthy donors living in Shenzhen, Guangdong province, China (GPS positioning of the samples collection site is 37°35’37”N/114°15’32”E) and preserved refrigerated and anaerobically until processed. The collection of the samples was approved by the Institutional Review Board on Bioethics and Biosafety of BGI under number BGI-IRB17005-T1. All protocols were in compliance with the Declaration of Helsinki and explicit informed consent was obtained from the participants. 1 g of faecal sample was diluted with 0.1 M PBS (pH 7, supplemented with 0.5% cysteine) and spread onto modified peptone-yeast extract-glucose (MPYG, supplemented with 5g/L sodium acetate in DSMZ 104 medium) agar plates in an anaerobic box (Bactron Anaerobic Chamber, Bactron□-2, shellab, USA). The plates were incubated at 37°C under anaerobic conditions (90% N_2_, 5% CO_2_ and 5% H_2_, v/v) for 3 to 5 days. Single colonies were randomly picked and purified by repetitive subculturing on the new plates containing the same medium and incubated under the same conditions as described above. Among the pure cultures, two isolates, designated as AF52-21^T^ and CM04-06^T^, respectively, were obtained and subsequently maintained in 20% (v/v) glycerol and frozen at −80°C.

### Phenotypic characterization

The morphological characteristics of strains AF52-21^T^ and CM04-06^T^ were performed on cultures grown on MPYG medium at 37°C. Bacterial cell shape was examined by phase contrast microscopy (Olympus BX51, Japan) during the exponential phase of growth. Cell motility was examined using semi-solid MPYG medium containing 0.5% agar^41^. The Gram reaction was carried out using a Gram-staining kit (Solarbio, China). Spore formation and presence of flagella were determined by staining using spore stain kit and flagella stain kit supplied by Solarbio (China) following the manufacturer’s instructions. Colony morphology was observed following growth of the cultures on PYG agar for 2 days at 37°C. Optimal temperature for growth was determined using growth in MPYG medium at 4, 10, 20, 25, 30, 35, 37, 45 and 50°C for 7 days. The pH range for growth was also measured in MPYG medium covering the range of pH 3.0–10.0 (at interval of 0.5 pH units) at 37°C for 7 days. Growth at various NaCl concentrations (0-6%, in increments of 1.0%) was performed for determining tolerance to NaCl. Bile tolerance was measured at different bile salt concentrations (0-5%, at an interval of 1.0%) in the MPYG medium. Catalase activity was assessed by gas formation after dropping the fresh cells in 3% H_2_O_2_ solution. Biochemical properties, including utilization of substrates, acid production from carbohydrates, enzyme activities, hydrolytic activities, were determined using the API 20A, API 50CHL and API ZYM systems ((bioMérieux Inc., Marcy-l’Étoile, France) according to the manufacturer’s instructions with modification by adding sodium acetate at concentration of 0.5% in all tests. The reference type strain was tested under the same condition with strains AF52-21^T^ and CM04-06^T^.

### Chemotaxonomic characteristics

Chemotaxonomic features were investigated by analyses of cellular fatty acids. Biomasses of strains AF52-21^T^ and CM04-06^T^ were harvested from cells growing in MPYG at 37°C for 2 days. Whole cell fatty acid methyl esters (FAMEs) were extracted, separated and identified according to the MIDI Microbial Identifications System and performed by CGMGG (China General Microbiological Culture Collection Center, Beijing, China) identification service.

### Fermentation products analysis

For analysis the metabolic end products from glucose fermentation, including SCFAs and organic acids, cells were cultured in MPYG broth at 37°C for 2 days. Supernatant harvested from the cultures centrifuged at 10000g for 10min was used for determining SCFAs and organic acids. SCFAs detection was performed using a gas chromatograph (GC-7890B, Agilent) equipped with a flame ionization detector (FID) and capillary column packed with Agilent 19091N-133HP-INNOWax porapak HP-INNOWax (30m × 0.25mm × 0.25um). Organic acids were analysed by equipping capillary column packed with Agilent 122-5532G DB-5ms (40m × 0.25mm × 0.25um).

### PCR of bacterial 16S rRNA genes and phylogenetic analysis

Total genomic DNA of strains AF52-21^T^ and CM04-06^T^ were extracted using the standard phenol:chloroform method as described by Cheng and Jiang^42^. The complete 16S rRNA genes were amplified and sequenced according to the method previously described^43^. Primers used for amplification of 16S rRNA genes were 27f (5’-AGAGTTTGATCATGGCTCAG-3’) and 1492r (5’-TAGGGTTACCTTGTTACGACTT-3’). The obtained 16S rRNA gene sequences of strains AF52-21^T^ and CM04-06^T^ were compared with the sequences of type strains retrieved from the EzBioCloud database (https://www.ezbiocloud.net/)44 using the BLAST program to determine the nearest phylogenetic neighbours and 16S rRNA gene sequence similarity values. Phylogenetic trees were reconstructed by using the neighbour-joining method^45^, maximum-likelihood method^46^ and minimum-evolution method^47^ with the MEGA X program package^48^, after Clustal W multiple alignment of the sequences. Robustness of the phylogenetic trees was evaluated by using the bootstrap resampling method (1000 resamplings) of Felsenstein^49^.

### Genome sequencing, assembly, and annotation of isolates

For genome sequences of strains AF52-21^T^ and CM04-06^T^, genome DNA was prepared following the method described above. The draft genome was sequenced on an Ion Proton Technology (Life Technologies) platform at BGI-Shenzhen (Shenzhen, China) after constructing a paired-end DNA library with insert size of 500 bp. The resulting reads were assembled using the SOAPdenovo 2 package^50^. CheckM (v1.1.2) was used to estimate genome completeness and contamination^51^. Genome assemblies were visualized using CGView Server^52^ (http://stothard.afns.ualberta.ca/cgview_server/index.html). Annotation of the assembled genome was performed using the Rapid Annotation Using Subsystem Technology (RAST) server^53^ and COG database^54^. The G+C content in genomic DNA was calculated from the whole genome sequence. The genes in known pathways from acetyl-CoA to butyrate were annotated by BLAST (evalue=1e-5, identity≥60%, coverage≥90%)^33^. A search for prophages was performed by PHAST (http://phast.wishartlab.com/)55. Antibiotic resistance was analysed using the CARD database^56^.

### Average nucleotide identities

Genome relatedness was investigated by calculating average nucleotide identity (ANI)^57^, with a value of 95-96% proposed for delineating bacterial species, corresponding to the traditional 70% DNA–DNA reassociation standard^58,59^. The ANI values between strains AF52-21^T^, CM04-06^T^, and closely related species were determined using the FastANI^60^.

## Acknowledgment

This work was supported by grants from National Key Research and Development Program of China (No. 2018YFC1313801) and Natural Science Foundation of Guangdong Province, China (No. 2019B020230001). We also thank the colleagues at BGI-Shenzhen for sample collection, and discussions, and China National Genebank (CNGB) Shenzhen for DNA extraction, library construction, and sequencing.

## Author contributions

Conceived and designed the experiments:Y.Z. and L.X. Performed the experiments: Y.Z., W.X., and Y.D. Analyzed the data: Y.Z., L.X., and X.L. Contributed reagents/materials/analysis tools: Y.Z., W.X.. and Y.D. Wrote the paper: Y.Z. and X.L. Revised the paper: K.K.

## Data Availability Statement

The GenBank/EMBL/DDBJ accession numbers for the 16S rRNA gene sequences determined in this study are: AF52-21^T^ (KX146426) and CM04-06^T^ (KX150462). The data of draft genome sequences have been deposited into CNGB Sequence Archive (CNSA) ^61^ of China National GeneBank DataBase (CNGBdb) ^62^ with accession number CNA0017730 and CNA0017731 for strains AF52-21^T^ and CM04-06^T^, respectively.

## Supplementary Material

**Supplementary Table S1. Number of genes associated with general COG functional categories in the genome of *F. butyricigenerans* AF52-21**^**T**^ **and *F. longum* CM04-06**^**T**^.

**Supplementary Table S2. The specific genes/protein related to biosynthesis of DAP, polar lipids, polyamines and lipoteichoic and teichoic acids and their positions in the genome in comparation of strains AF52-21**^**T**^, **CM04-06** ^**T**^ **and related organism, ATCC 27768**^**T**^ **identified by Rapid Annotation Subsystem Technology (RAST)**.

**Supplementary Figure S1. Maximum-likelihood phylogenetic tree based on 16S rRNA gene sequences showing the phylogenetic relationships of strains AF52-21**^**T**^, **CM04-06**^**T**^ **and the representatives of related taxa**. *Clostridium butyricum* DSM 10702^T^ (AQQF01000149) was used as an out-group. Bootstrap values based on 1000 replications higher than 70% are shown at the branching points. Bar, substitutions per nucleotide position.

**Supplementary Figure S2. Minimum-evolution phylogenetic tree based on 16S rRNA gene sequences showing the phylogenetic relationships of strains AF52-21**^**T**^, **CM04-06**^**T**^ **and the representatives of related taxa**. *Clostridium butyricum* DSM 10702^T^ (AQQF01000149) was used as an out-group. Bootstrap values based on 1000 replications higher than 70% are shown at the branching points. Bar, substitutions per nucleotide position.

**Supplementary Figure S3. Certification. Deposit certification of AF52-21**^**T**^ **in CGMCC**.

**Supplementary Figure S4. Certification. Deposit certification of AF52-21**^**T**^ **in DSMZ**.

**Supplementary Figure S5. Certification. Deposit certification of CM04-06**^**T**^ **in CGMCC**.

**Supplementary Figure S6. Certification. Deposit certification of CM04-06**^**T**^ **in DSMZ**.

