## Supplementary Figure S6 for "Characterization and description of *Faecalibacterium butyricigenerans* sp. nov. and *F. longum* sp. nov., isolated from human faeces"

Leibniz-Institut
DSMZ-Deutsche Sammlung von
Mikroorganismen und Zellkulturen GmbH

DSMZ

**Confirmation of the availability of a strain for the purpose of valid publication of a new name according to the Bacteriological Code**

The following information is confidential and serves only to allow the International Journal of Systematic and Evolutionary Microbiology to confirm that a strain has been deposited and will be available from the DSMZ in accordance with the Rules of the Bacteriological Code (1990 revision) as revised by the ICSP at the plenary sessions in Sydney and Paris.

Strain **Faecalibacterium longum CM04-06** has been deposited in the DSMZ under the number

**DSM 103432**

This strain is available in the publicly accessible section of the DSMZ and restrictions have not been placed on access to information concerning the presence of this strain in the DSMZ. It will be included in published and online catalogues after publication of this number by the authors.

This strain has been checked for viability in the DSMZ and is stored using one of the standard methods used in the DSMZ. The depositor of this strain has also carried out a "depositor's check" and confirmed the identity of the strain held under this DSM number.

**!! The DSMZ is not responsible for differences between the properties of the strain deposited in the DSMZ and properties given in the literature/databases !!**

**It is the sole responsibility of the depositor to ensure that type strains deposited in the DSMZ conform to the requirements of the appropriate Rules governing prokaryotes nomenclature and the deposition of type strains (Rules 18a, 27, & 30 of the ICNB/ICNP, including changes made at plenary sessions of the JC/ICSP)**.

Dr. R. Pukall, Curator Gram-positive Bacteria

Geschäftsführer/
Managing Director:
Prof. Dr. Jörg Overmann
Aufsichtsratsvorsitzender/Head of
Supervisory Board: RD Dr. David Schnieders

Braunschweigische Landessparkasse
(NORD/LB) Kto.-Nr./Account: 2 039 220
BLZ/Bank Code: 250 500 00
IBAN DE22 2505 0000 0002 0392 20
SWIFT (BIC) NOLADE 2 H

Handelsregister/
Commercial Register:
Amtsgericht Braunschweig
HRB 2570
Steuer-Nr. 13/200/24030

DAkkS
Deutsche
Akkreditierungsstelle
D-RM-18324-01-00
