## Supplementary figures and images for "Characterization and description of *Faecalibacterium butyricigenerans* sp. nov. and *F. longum* sp. nov., isolated from human faeces"

### Supplementary Figure S1

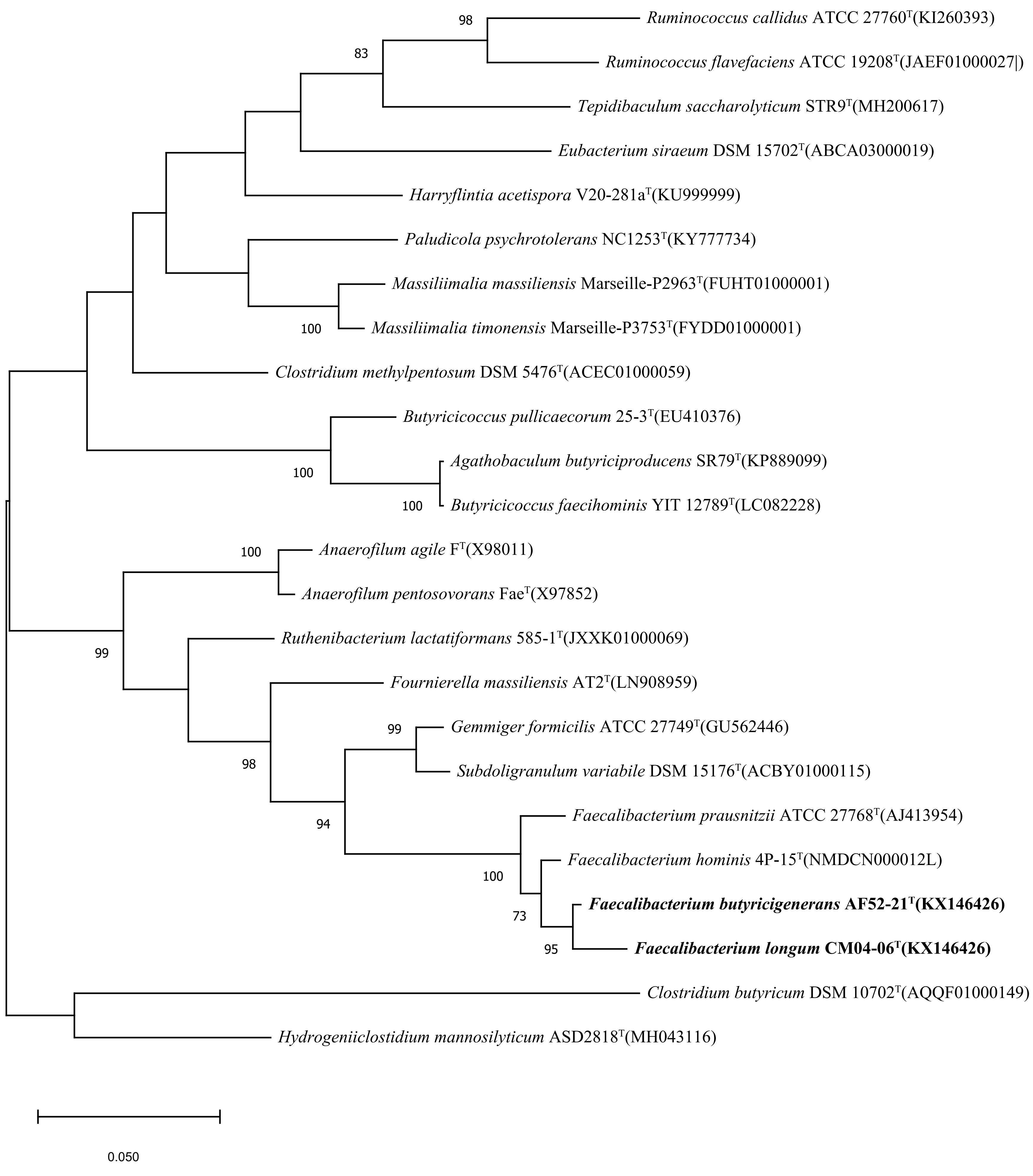

### Supplementary Figure S2

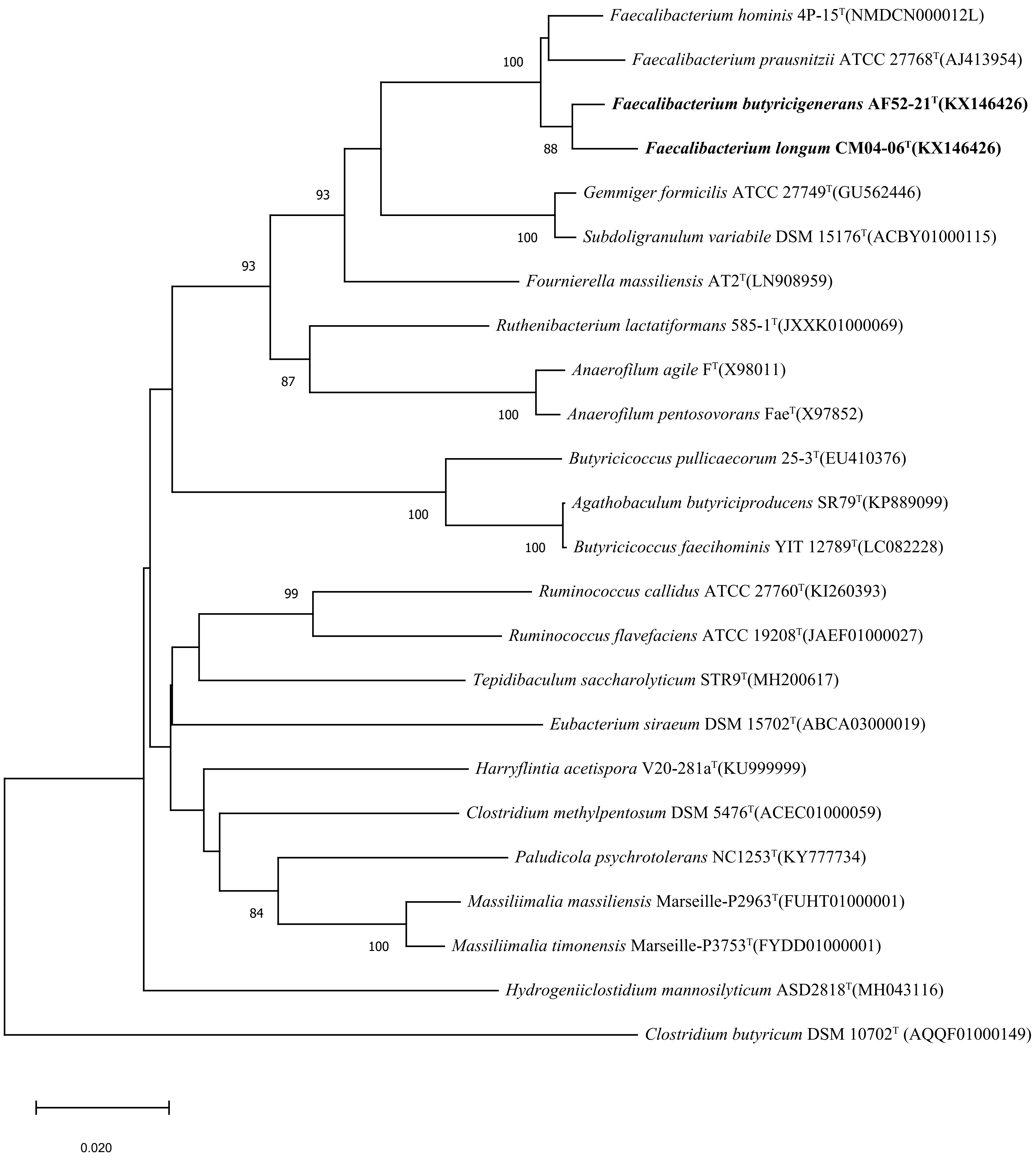
