## Supplementary Figure S3 for "Characterization and description of *Faecalibacterium butyricigenerans* sp. nov. and *F. longum* sp. nov., isolated from human faeces"

### 中国微生物菌种保藏管理委员会普通微生物中心
### China General Microbiological Culture Collection Center (CGMCC)

Address: Institute of Microbiology, Chinese Academy of Sciences, Datun Road, Chaoyang District, Beijing 100101, China

#### 受理通知书
#### NOTIFICATION OF RECEIPT

**CGMCC 1.5206**

1. Name and address of the depositor or agent

薛文斌 Wen-Bin XUE
BGI-Shenzhen
Main Building, Beishan Industrial Zone, Yantian District, Shenzhen 518083, China

2. Strain reference given by depositor

**AF52-21**

3. Deposited microorganisms appended

☐ Scientific description

■ Proposed taxonomic name

***Faecalibacterium butyricigenerans***

4. The deposited microorganism has been received and numbered as CGMCC 1.5206 on January, 2016. The strain has been checked for viability in the CGMCC and is stored using one of the standard methods used in the CGMCC.

5. This strain is available in the public accessible section of the CGMCC and restrictions have not been placed on access. It will be included in the published and online catalogue after publication of this number by the authors.

Signature of Head of CGMCC Yu-Guang ZHOU

Date June 13, 2016

中国微生物菌种保藏管理委员会普通微生物中心
